# A Narrative Systematic Review of Coaching Interventions to Improve Dyslexia at Work

**DOI:** 10.1101/342584

**Authors:** Nancy Doyle, Almuth McDowall

## Abstract

Although dyslexia affects 5-8% of the workforce this developmental disorder is insufficiently researched within adult psychological research. Dyslexia confers legal protections wherein employers must provide ‘accommodations’ to support work performance, including coaching activities. Implementation of accommodations has moved forward without an evidence-base to date. The present systematic review investigates to what extent face-to-face learning interventions improve the working memory capacity and self-efficacy of adults with dyslexia guided by a realist, narrative framework. We extracted 25 studies from eleven countries, prioritizing relevant learning intervention protocols above population context, synthesizing the delivery context and impact on mechanisms of working memory (N=501) and self-efficacy (N=1211) also contextual, work-related outcomes such as comprehension. Though limited by inadequate intervention descriptions in primary papers, a narrative synthesis determined that learning interventions compliant with Social Cognitive Learning Theory elicited reliable outcomes for not only self-efficacy, as would be expected, but also improved working memory and workplace contextualized variables. Development of metacognition, stress management and fidelity to Goal Setting Theory were also inferred as valuable intervention features. Implications include the need for population-specific evaluation of the emergent conceptual framework to further our understanding of disability accommodations, and questioning the ecological validity of working memory as an intervening mechanism.

## Introduction

Developmental disorders including dyslexia are common in adult working populations, affecting about 5-8% of the workforce (Rice & Brooks, 2004) but our understanding of how affected individuals can be supported to facilitate career success and well-being remains under-developed. In the U.S. the Disabilities Act (U.S. Equal Employment Opportunities Commission, 2008) defines as a physical and/or mental impairment that has limiting effects on any life activity; the corresponding UK definition refers to physical or mental impairments that have a ‘substantial’ and ‘long-term’ negative effect on people’s ability to engage in normal daily activities (United Kingdom Parliament, 2010). Both Acts require employers to make ‘accommodations’ (US terminology, which we adopt for the purposes of this paper) or ‘adjustments’ (UK terminology) to ensure that workers are not disadvantaged. Dyslexia is subject to relevant employment legislation in many advanced economies including the UK, Canada and the USA (de Beer, Engels, Heerkens, & van der Klink, 2014; Gerber, 2012; Gerber, Batalo, & Achola, 2012). In the UK alone, over 3000 individuals per year each receive approximately $1000 of public funding to access support, including technology and coaching (Gifford, 2011; Melvill, Stevens, & Vaid, 2015)

Whilst workplace coaching activities are habitually dispensed as disability accommodations for dyslexic employees (Bewley & George, 2016; Doyle & Mcdowall, 2015; Melvill et al., 2015) there is a gap in psychological research with regards to theoretical framing for reviewing the purported effectiveness. It is the purpose of the current paper to (1) set out a conceptual context for coaching disability accommodations, including psychosocial factors and theories relevant to dyslexia, and then (2) formulate a clear research question to (3) guide a systematic review on the evidence for the theoretical framing and effectiveness of learning interventions, in order to (4) inform dyslexia coaching practice.

### Psychosocial Factors in Dyslexia

Dyslexia is associated with an increased vulnerability to criminal activity (Snowling, Adams, Bowyer-Crane, & Tobin, 2000), higher unemployment (Jensen, Lindgren, Andersson, Ingvar, & Levander, 2000), failure to achieve potential post-education (Holliday, Koller, & Thomas, 1999) and impaired workplace participation (defined by de Beer et al., 2014, p. 4, as “work content, work circumstances, terms of employment and relationships at work”;WHO, 2001). To illustrate, only 1% of corporate managers are dyslexic (Logan, 2009) compared with a population norm of 10% (Snowling, 2010) demonstrating an unequal pattern in career achievement. The critical nature of dyslexia accommodations for social inclusion and equality (Kim, Sally, & Joseph, 2002; O’Brien & Ellegood, 1996; von Schrader, Malzer, & Bruyère, 2014) creates a moral imperative to ensure accommodations are well researched. However, the balance of dyslexia research is strongly weighted toward children and educational needs only (Doyle & Mcdowall, 2015).

## Dyslexia Theory

We provide a brief overview regarding the theoretical inconsistencies in the dyslexia literature particularly given that available evidence is concentrated in the educational domain (Kirby, Sugden, Beveridge, Edwards, & Edwards, 2008; Lyon, Shaywitz, & Shaywitz, 2003; Mortimer & Crozier, 2006; Richardson & Wydell, 2003) to inform our review. Understanding of adult symptomatology remains underdeveloped, not the least because most research relies heavily on correlational data and designs (Burden, 2008).

Historically, dyslexia was thought to be a visual processing disorder (“word blindness”; Orton, 1937). A deficit in phonological processing is the current dominant theory in developmental and educational research (Bishop, Snowling, & Blakemore, 2007; Shaywitz, 1998; Vellutino, Fletcher, Snowling, & Scanlon, 2004) though rapid-naming (visual recognition of words at speed), which was originally proposed as a causal factor by Denckla & Rudel (1976) is still routinely tested in diagnosis (Grant, 2009; McLoughlin & Doyle, 2013; McLoughlin & Leather, 2013) and therefore in practice, the double-deficit hypothesis (Wolf & Bowers, 1999 - meaning both difficulties in rapid naming and the accurate decoding of sounds) is still used to define and determine individual cases. Further to phonological deficit theory, neuropsychological researchers have highlighted the phonological short-term component of ‘working memory’ (our capcity to hold information in our attention long enough to manipulate it, Baddeley, 2000) as the primary neural mechanism leading to phonological processing difficulties, which in turn result in delayed literacy acquisition (Jeffries & Everatt, 2004; Smith-Spark, Fisk, Fawcett, & Nicolson, 2003; Swanson & Siegel, 2001; Torgesen, 2001). Other lines of research include exploring different long-term memory-based hypotheses, such as the cerebellar deficit theory, which implicates a lack of automatization and issues with balance/motor control (Nicolson, Fawcett, & Dean, 2001). Research suggests that the neuropsychological structure of dyslexia may vary according to the language that one speaks (Opitz, Schneiders, Krick, & Mecklinger, 2014; Siok, Spinks, Jin, & Tan, 2009), and some researchers remain unconvinced that there are any specific neuropsychological elements that are sufficiently distinct to differentiate dyslexia from general poor reading skills (Elliot & Grigorenko, 2014). The variation between, and number of, probable causal psychological mechanisms leaves an absence of theoretical direction for evaluative research. De Beer et al.’s (2014) systematic review of factors influencing work participation for dyslexia suggested that coaching as a disability accommodation could have many potential psychosocial outcomes, including cognitive, emotional, behavioral, social and environmental factors.

Given such inconclusive extant literature and the imbalance of research toward childhood literacy attainment, it was first necessary to conduct a scoping process, through consultation with an expert panel and reviewing survey data prior to our actual search, in order to refine the conceptual framework for the review, in accordance with guidelines on SR methodology (Denyer & Tranfield, 2009).

## Scoping the Research Question

### Expert panel consultation and user data review

We consulted a virtual panel of 14 internationally recognized experts across research and practice, including academically published educational, occupational and clinical psychologists, educational neuropsychological researchers and one dyslexia specialist from the charitable sector, to inform our protocol. Through structured telephone interviews, we questioned (1) their understanding of relevant psychological variables for adults with dyslexia; (2) potentially under-researched but urgent topics, as well as; (3) direct questions about pertinent literature. We then conducted thematic mapping of their responses. Regarding symptomatology, 51% of responses made direct references to Working Memory (WM) or indicators of WM-related behaviors. Psychosocial difficulty was the second most frequent response (27%) and the need to support Self-Efficacy (SE) beliefs was highlighted. WM is an executive cognitive function that is considered a component of crystallized intelligence (Baddeley, 2000). SE refers to the development of our perceived ability to perform a given task and is a central tenet of Social Cognitive Learning Theory (SCLT; Bandura, 1986) and has been raised in contemporary research as particularly problematic for dyslexic adults (Doyle & McDowall, 2015; Leather et al., 2011; Nalavany, Logan, & Carawan, 2017). The entire panel stressed the need for evaluations of current practice interventions, specifically coaching, and contributed to the search terms.

To embed the perspective of the end user in the SR scoping process (Riddick, 2001; Shakespeare & Watson, 1997) we reviewed existing survey data from 81 working dyslexic adults (Doyle & Cleaver, 2015) which documented the focal topic of any disability accommodation requests, the most popular being memory support, highlighted by 71% of respondents. In conclusion, congruence between stakeholders, expert panelists and contemporary research suggested that WM and SE were the most important psychological variables to target in providing support for adults with dyslexia. To ensure that targeting these variables would be justifiable in the context of disability accommodation, we now briefly outline the relationship between both variables and work performance, as well as explore the use of coaching as a psychological intervention.

## Key Terminology

### Working Memory and Work Performance

WM is linked to complex cognitive reasoning (Ariës, Groot, & van den Brink, 2014; Conway, Kane, & Al, 2005; Hofmann, Schmeichel, & Baddeley, 2012; Klingberg, 2009; Swanson & Siegel, 2001) and is implicated in a range of effective work-related behaviors, such as self-regulation (Wolf & Kaplan, 2008), time management (Mantyla & Carelli, 2006) and management of complex environments (Thorell, Lindqvist, Nutley, Bohlin, & Klingberg, 2009). The clear association between WM and work performance, vindicates its inclusion in disability accommodation activities, yet leaving the question of *how* WM could be improved via coaching unanswered. We noted at this stage the more developed and growing body of literature on improving WM through adaptive computerized training and the dissonant results between successful ‘near transfer’ of WM skills and less successful ‘far transfer’ of higher cognitive reasoning skills akin to a contextualized work performance. Systematic reviews of computerized interventions demonstrate that when WM is targeted specifically it improves, but these improvements fail to translate to wider successes across a wide range of client groups (children with reading disabilties: Dunning, Holmes, & Gathercole, 2013; clinical neuropsychological rehabilitation and healthly populations: Melby-Lervåg, Redick, & Hulme, 2016; Weicker & Thöne-otto, 2015). For this reason, higher cognitive skills, contextualized skills and specific work performance related skills were specifically extracted and analyzed in the synthesis, to consider if coaching interventions experienced the same comparatively weaker effect with contextual measures.

### Self-Efficacy and Work Performance

Functioning and performance in the employment context are contingent on supportive interactions with others and on positive self-belief (de Beer et al., 2014; WHO, 2001). A previous meta-analysis by Stajkovic and Luthans (1988) showed that SE, similar to WM, has a strong relationship with work performance in the general population, and in particular, high SE has been shown to mediate the impact of poor workplace outcomes in people with dyslexia (Gerber et al., 2012; Leather et al., 2011; Werner, 1993). To explore the viability of coaching as a disability adjustment for dyslexia, coaching interventions targeting SE in the workplace were thus justified as the second focus of our review.

### Coaching for dyslexia

Following the scoping phase we reviewed the definition and nature of coaching used in this very specific context. The form of coaching commonly used in disability accommodation (Bewley & George, 2016; Doyle, Cleaver, & Rossiter, 2016) is not a straightforward continuation of the various better-researched coaching/tuition interventions provided in education (Mortimer & Crozier, 2006; Rose, 2009). Coaching to support people with dyslexia in the workplace does not focus on literacy attainment but rather on outcomes more commonly associated with general workplace coaching such as time management and organizational skills (Doyle & Mcdowall, 2015; McLoughlin & Leather, 2013). Coaching is aimed at cognitive, behavioral and emotional changes rather than knowledge transfer, and thus lends itself to a more dialectic than didactic pedagogy. Coaching psychology literature defines workplace coaching as a developmental activity for individuals and organizations (Grant, Passmore, Cavanagh, & Parker, 2010) aiming to unlock an individual’s potential (Whitmore, 1992) to ultimately improve workplace productivity and job performance. Workplace coaching objectives are thus more aligned to disability accommodation than literacy acquisition. Specifically, McLoughlin and Leather (2013, p.43) describe workplace dyslexia coaching as an “androgogical approach” that relies on the metacognitive experience of dyslexic adults (Leather et al., 2011) but they highlight that deviations from this style are common in practice. Some ‘coaches’ resort to literacy tuition, training and knowledge transfer, very similar to educational interventions (Doyle & Mcdowall, 2015). The extent to which interventions adhered to dialectic, rather than didactic, principles was thus explicitly considered in the sampling and synthesis of primary studies. We note the divergence in learner experience between a coaching psychology intervention and computerized working memory training; a self-directed, yet conversational and social learning protocol compared to a solitary, technology-based exercise, practicing similar tasks repetitively.

## Summary of the Review Aims

Coaching as a disability support for dyslexia at work has lacked a psychological evidence base thus far, despite current popularity in practice, with implications for a vulnerable client group consisting of up to 10% of the working population (IDA, 2002; J. R. Kirby, Silvestri, Allingham, Parrila, & La Fave, 2008; Rice & Brooks, 2004). The purpose of this review was therefore to address and scrutinize existing treatment studies through a focus on potential WM and SE improvements to investigate psychological mechanisms for relevant coaching interventions that could be effective for improving dyslexic difficulty in the workplace. We specifically drew out the WM results and compared these to adaptive, computerized training of WM which has received considerably more research interest to date (Melby-Lervåg et al., 2016; Weicker & Thöne-otto, 2015).

## Method: Review Protocol

### Narrative Synthesis Using the CIMO Framework

Given the likely paucity of primary studies and heterogeneity in methods, we used a ‘Realist Synthesis’ approach which, in contrast to traditional reviews of medical evidence (The Cochrane Collaboration, 2008), develops a narrative theoretical explanation of “how and why programs work” (Pawson, 2006, p. 74). We built on the context, intervention, mechanisms and outcomes (CIMO) framework recommended by Rousseau, Manning, & Denyer (2008), and focused on the two core psychological variables derived from the exploratory scoping phase to create a hypothetical framework for evaluation: (1) face-to-face intervention evaluations, (2a) WM and (2b) SE as target intervening variables for dyslexic adults in occupational contexts. Table 1 depicts our interpretation of the CIMO framework,

**Table 1.** Interpretation of the CIMO framework for the current realist synthesis-

| Realist synthesis component | Explanation (Denyer & Tranfield, 2009) | Relevance to the present review |
| --- | --- | --- |
| Context | Individuals of interest.<br>Interpersonal relationships of interest.<br>Institutional setting of interest.<br>Aspects of wider infrastructure of interest. | Realistic approach to lack of context specific studies for WM<br>Include adults and children.<br>Include both men and women.<br>Include all nationalities.<br>Consider educational, health and occupational contexts. |
| Interventions | The intervention of interest. | Exclude medical- or technology-based |
|  |  | memory training interventions.<br>Include interventions described as ‘learning’ or ‘coaching’.<br>Include interventions delivered via face-to-face dialectic pedagogy.<br>Exclude didactic-type interventions.<br>Controlled studies only. |
| Mechanism | Mechanisms of interest.<br>Explanation of how the interventions act within the context to lead to the outcome.<br>Understanding of how mechanisms are activated or not activated in different contexts. | Unusual to the CIMO structure, WM and SE were both considered mechanisms and outcomes in the search for studies, provided that the study included a standardized measure of either mechanism as a dependent variable.<br>Additional mechanisms (contributing to WM/SE improvement) and work-related outcomes, such as higher order reasoning abilities, were also considered in the extraction and synthesis. |
| Outcome | Relevant outcomes.<br>Measurement of outcome.<br>Primary and secondary outcomes. |  |

## Context and Intervention

The contextual search terms were broadened, a necessary concession to the lack of primary papers, allowing us to propose hypothetical principles, which can then be tested in further studies, rather than a direct match with dyslexic samples. For WM, all relevant adult populations (healthy or not, education- or work-based) were included with the exception of: (1) samples with serious age-related cognitive impairment (e.g., dementia) due to the multiple cognitive, social, health and clinical concerns of this group and; (2) child-based studies in which the interventions focused on the actions of the parents or teachers rather than the children themselves. As such, the following extraction digresses from the dyslexia literature to a more generalist psychological exploration of variables, and the potential impact for dyslexia will be addressed in the discussion.

Due to the aforementioned inconsistency in the definitions of coaching in the disability context, we broadened the initial search to include all face-to-face interventions, as defined by the absence of technology rather than the inclusion of coaching in particular; this differentiation allowed us to compare the WM extraction to systematic reviews of WM computerized training interventions. The resulting extracted studies were then analyzed specifically regarding the extent of their fidelity to a dialectic, recognized coaching definition, such as the one described above by McLoughlin and Leather (2013) and known to be successful in related fields such as ADHD (Parker & Boutelle, 2009; Richman, Rademacher, & Maitland, 2001).

## Mechanisms and Outcomes

Studies that did not include WM or SE as a clear mechanism or outcome were excluded, for example, when WM or SE was an independent rather than a dependent variable.

### Working memory

WM-focused studies were identified through inclusion of a published, standardized WM test as a dependent variable. Work-related outcomes were also extracted for consideration as secondary dependent variables, for example reading comprehension. Therefore, we incorporated studies that included WM as both a mechanism (process or intervening variable) and an outcome (dependent variable).

### Self-efficacy

SE-based studies were identified through the explicit use of the term, through reference to SCLT, and use of a validated SE scale.

### Research Question

Based on the above considerations, we seek to identify any indication for the potential effectiveness of habitual coaching accommodations which can be extrapolated from extant research. The primary, two-part question guiding our review was as follows: to what extent, and under what conditions, can face-to-face (C) learning interventions (I) improve WM (MO_1_) and SE (MO_2_)?

### Search Criteria

We mapped our search terms against the CIMO framework, as depicted in Table 2. The search used all relevant EBSCO-hosted databases and was concluded in June 2016, identifying 609 studies for WM and 414 for SE; data from these studies were recorded using a bespoke data extraction form. The terms used were as broad as possible and were cross-referenced with the expert panel to ensure that the terminology reflected the phrases commonly used in the numerous disciplines contributing to the review. As advocated by Denyer & Tranfield’s (2009) CIMO model we prioritized Intervention and Mechanism criteria through weighting.

**Table 2.** . Search terms

| **CIMO stage** | **Search terms** | **Search location**<br>**or stage** |
| --- | --- | --- |
| **Context** | Coaching OR training OR classroom OR<br>professional development OR intervention OR<br>activity OR learning OR face-to-face OR tuition<br>OR educat* | All Text |
| **Primary**<br>**context of**<br>**interest** | Dyslexi* OR adults OR 19+ | Filtering term<br>applied after the<br>initial search to<br>identify high<br>relevance<br>studies |
| **Interventions** | Learning OR metacognitiv* OR self-awareness<br>OR self-development OR synesthe* OR<br>synaesthe* OR instruct* Or knowledge OR<br>personal development | All Text |
| **Mechanism /**<br>**outcome 1** | Working memory OR executive function* OR<br>attention OR short-term memory OR cognition<br>OR metacogniti* OR time management OR self-regulation OR synesthe* OR synaesthe* OR<br>mental function* | Title / subject /<br>abstract /<br>keywords |
| **Mechanism /**<br>**outcome 2** | OR self-efficacy OR perceived self-efficacy OR<br>work efficacy OR self-efficacy belief OR social<br>cognitive learning theory OR social learning | Title / subject /<br>abstract /<br>keywords |
|  | theory OR self-esteem OR self-confidence OR<br>participation OR social interaction OR agency<br>OR career agency |  |

## Relevance and Quality Assessment

### Screening process

We screened by title, abstract (retaining 89 papers for WM and 84 for SE), removing any duplicates, and then full text (16 papers for WM and 38 for SE) and directly located primary studies by scanning the references of the extracted papers and book chapters (adding four papers). The expert panel was sent a full bibliography, and each member checked this list against their expert knowledge of relevant studies; three further papers were added following their recommendations. All remaining papers (23 for WM, 38 for SE) were subjected to an in depth relevance check using bespoke a priori criteria based on CIMO. A quality review was then conducted drawing on previously established quality criteria (Rojon, McDowall, & Saunders, 2011; full criteria available from the second author on request), adapted to include the management of bias in the respective study design, for example, use of a control group, blind or double-blind protocols and addressing of sampling errors.

Relevance for inclusion was cross-checked between the first and second authors for inter-rater agreement and any disagreements resolved by consensus through a second iteration involving joint examination of the primary papers. The main obstacle to assessing relevance was the persistent lack of detail in the papers regarding the nature of the intervention. For instance, in education research, the term ‘training’ was used interchangeably to refer to computer-guided adaptive practice and face-to-face learning. In ADHD research, many interventions considered themselves ‘psycho-social’ or ‘coaching’ but upon close inspection were actually interventions that targeted teachers’ or parents’ behaviors rather than coaching of the individual. Several authors were contacted to provide more information, but no responses were received.

For WM interventions we included seven studies of high quality (Alloway & Warner, 2008; Ariës et al., 2014; Chambers, Lo, & Allen, 2008; Craik et al., 2007; Miranda, Presentación, Siegenthaler, & Jara, 2013; Zeidan, Johnson, Diamond, David, & Goolkasian, 2010; Zylowska, Ackerman, Yang, Futrell, Horton, Hale, Pataki, & Smalley, 2008) and three of intermediate quality (Jha, Stanley, Kiyonaga, Wong, & Gelfand, 2010; Moro et al., 2012, 2015); excluding one study (Toll & Van Luit, 2013) for failing to include an appropriate WM measure post-intervention. A high quality paper (Ariës et al., 2014) was analyzed as two studies, as it included two data sets that measured the same outcomes but using separate samples and interventions.

For SE we retained fourteen studies, including eight studies of high quality (Franklin & Doran, 2009; McDowall & Butterworth, 2014; Mcdowall, Freeman, & Marshall, 2014; McGonagle, Beatty, & Joffe, 2014; Reif, De Vries, Petermann, & Görres, 2013; Tschannen-Moran & McMaster, 2009; Watt, Murphy, Pascoe, Scanlon, & Gan, 2011; Zwerver, Schellart, Anema, & Van Der Beek, 2013); and six of intermediate quality (Bell, Raczynski, & Horne, 2010; Engin & Cam, 2009; Reed, Kennett, Lewis, & Lund-Lucas, 2011; Stensrud, Gulbrandsen, Mjaaland, Skretting, & Finset, 2014; Style & Boniwell, 2010; Tsai et al., 2011). Two studies were excluded at this stage, due to lacking information on when post-intervention data were collected (Ogan-Bekiroglu & Aydeniz, 2013) and lacking an adequate baseline measure (Platt, 2011). Fig. 1 shows the number of papers included at each stage of the extraction.

**Figure 1.**
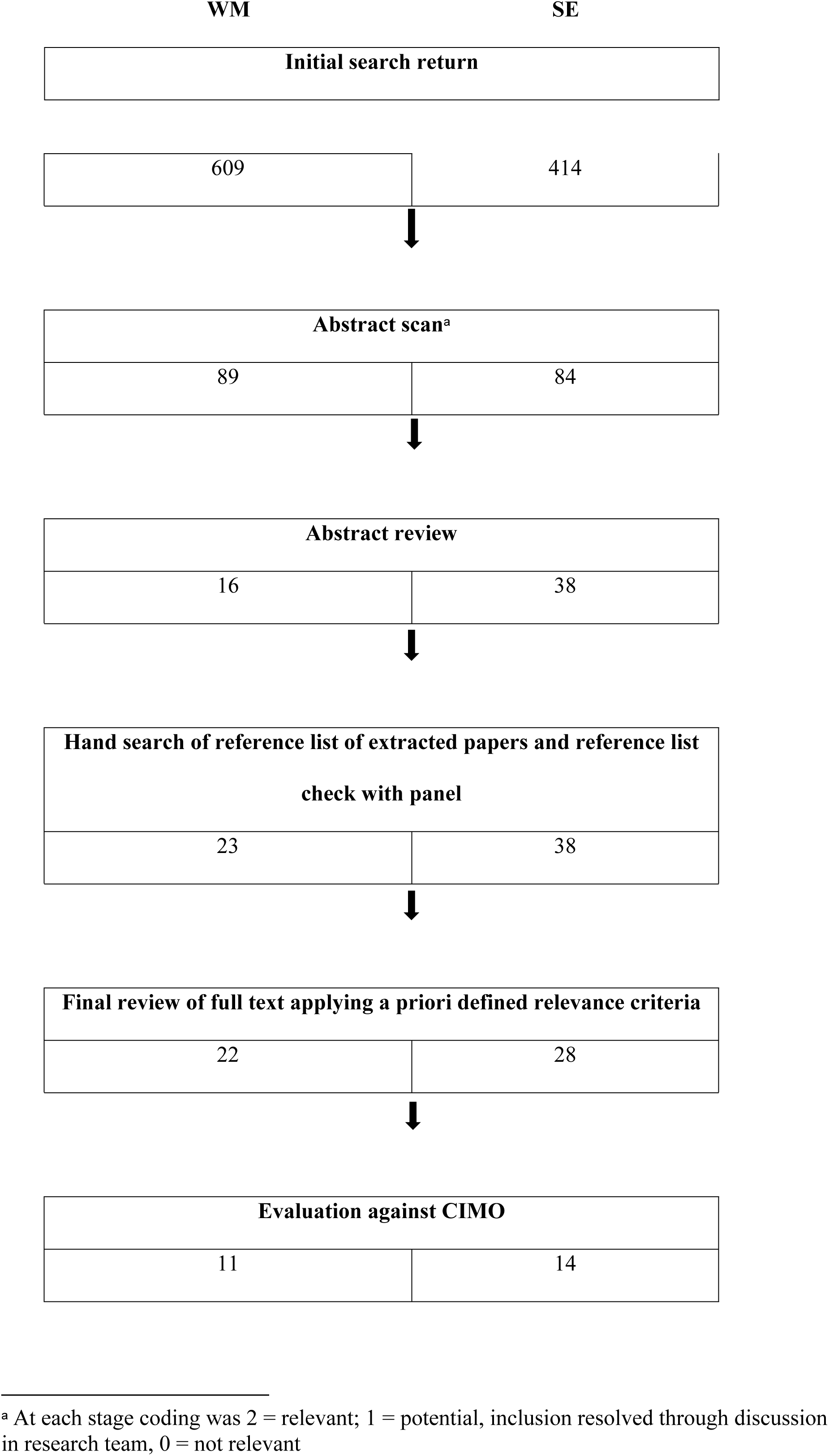
the initial sifting process and the number included at each stage.

## Results: Data Extraction and Synthesis

We synthesized the primary studies separately for WM and SE to compare the main elements in the CIMO framework, by (a) reviewing the effect sizes reported in each study (some of these were calculated by the primary author from provided means and standard deviations, shown in Table 3 and 4); followed by (b) grouping the studies according to outcome; and then finally (c) examining contexts and interactions to identify any common themes.

**Table 3.**
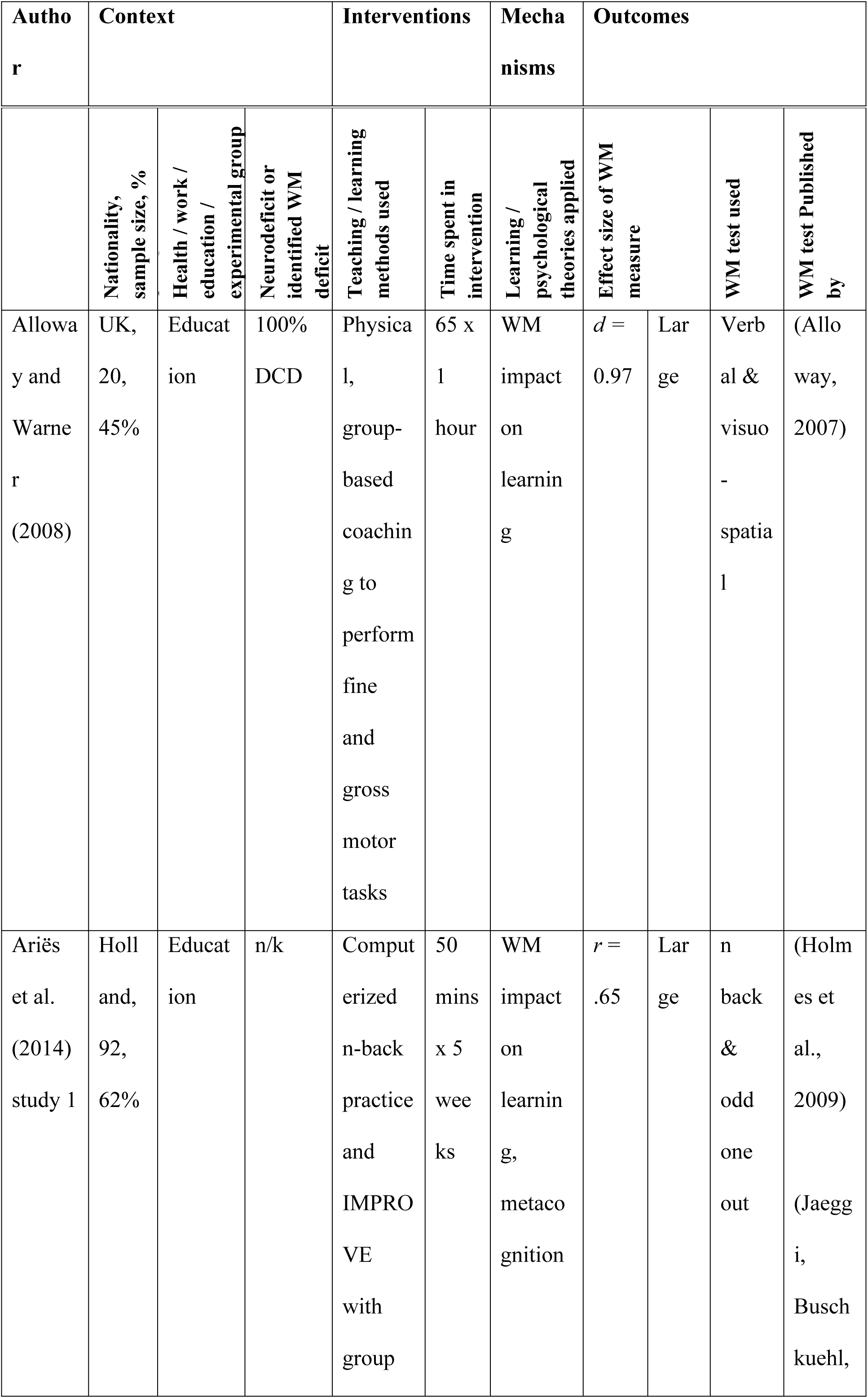
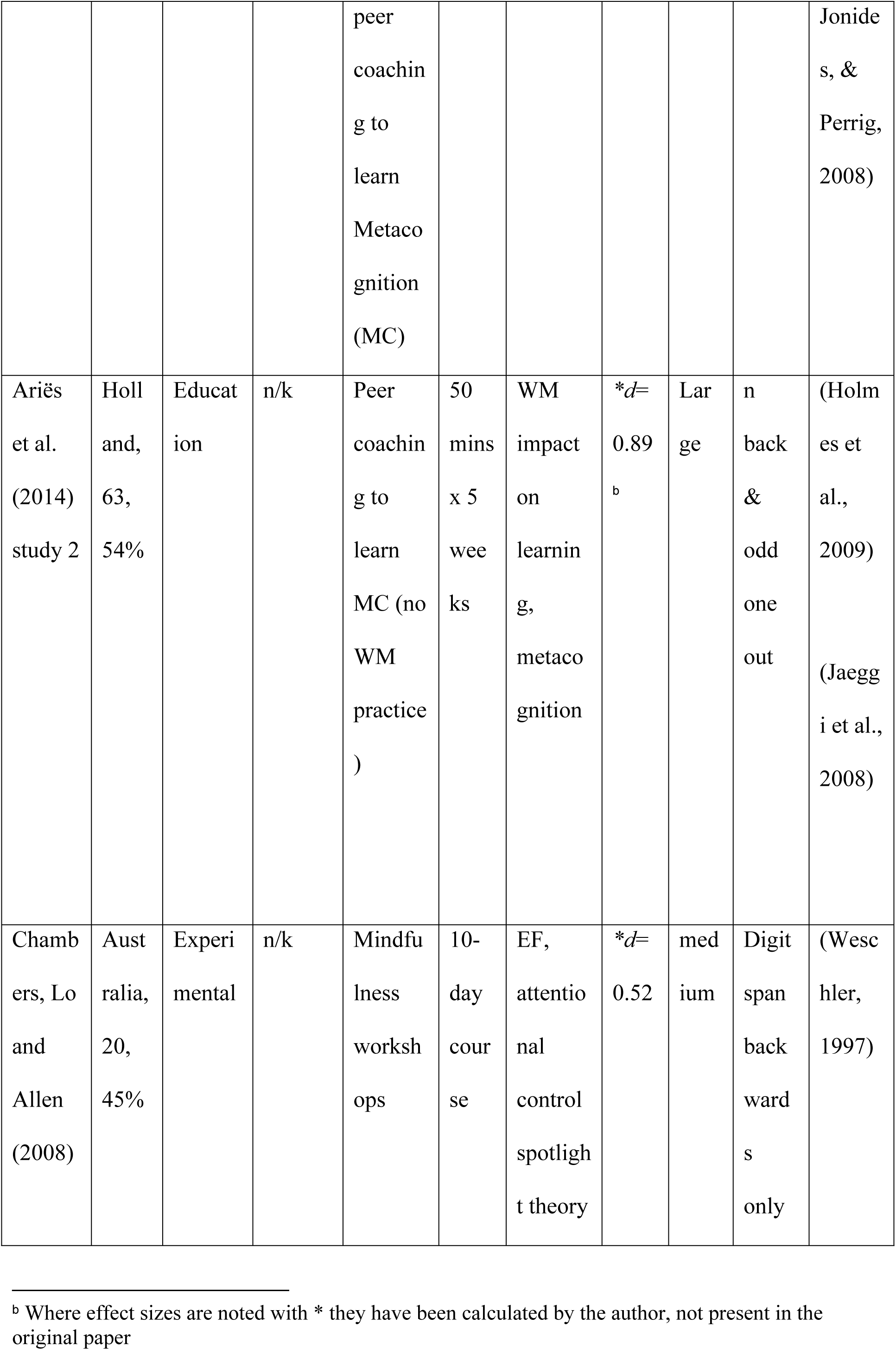
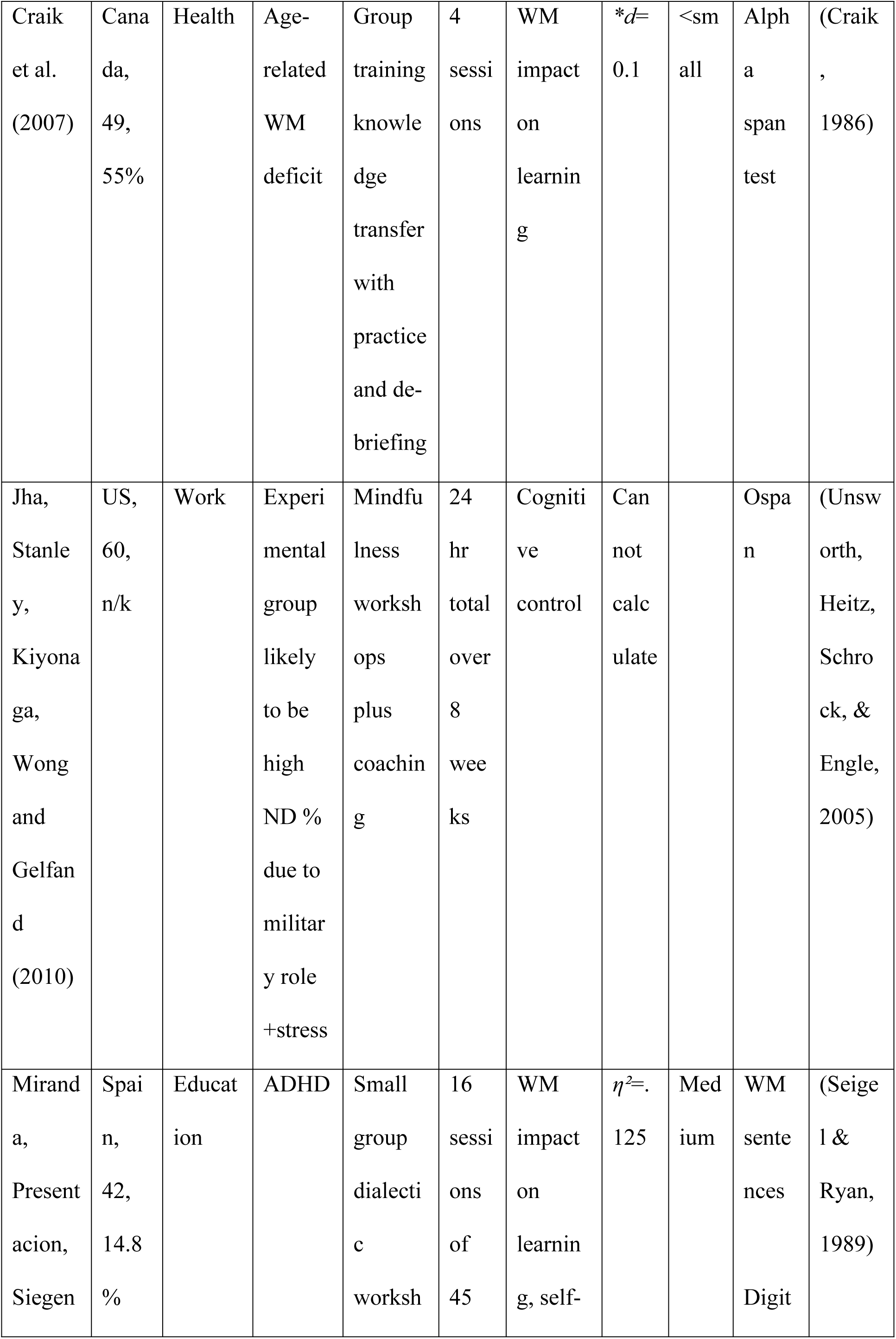
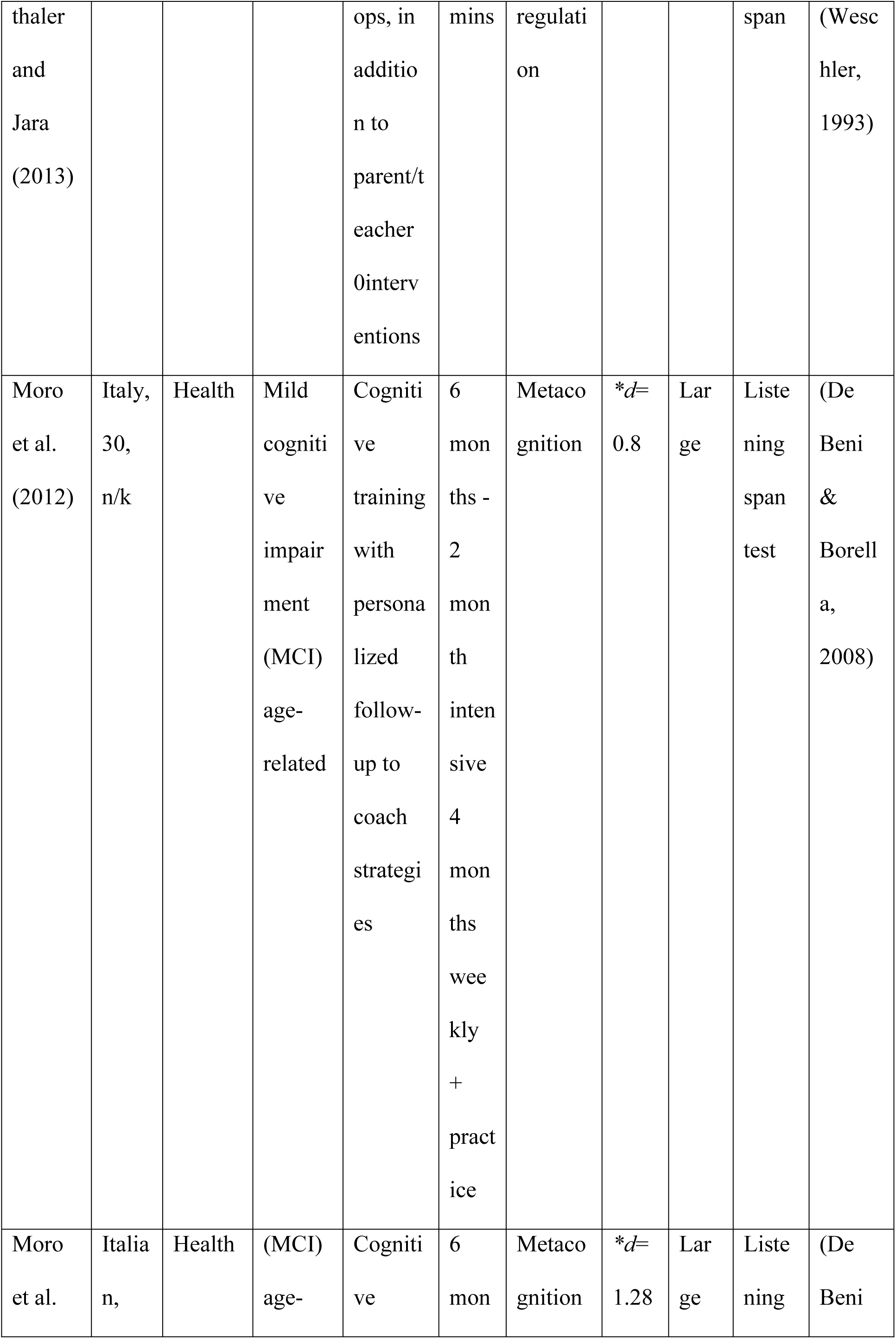
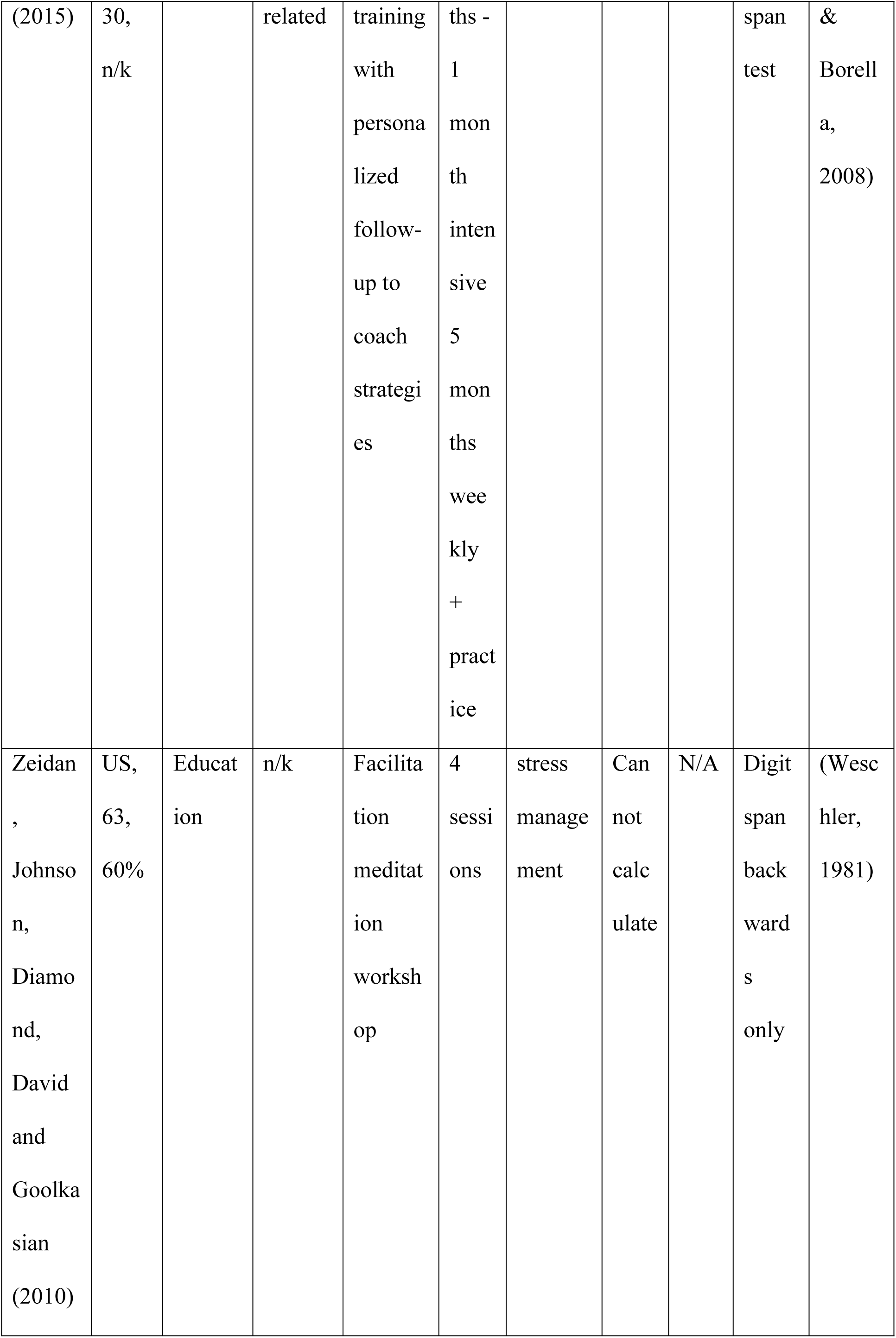
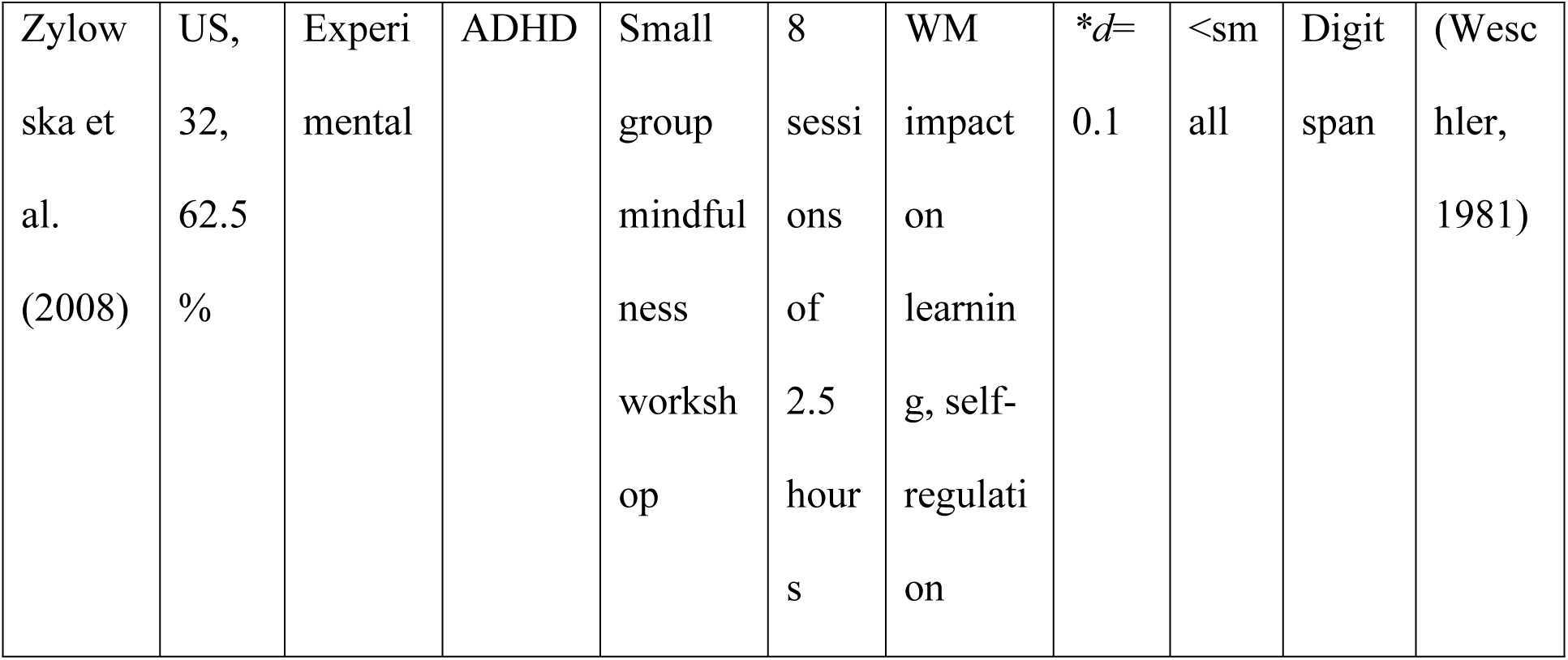
Data extracted from working memory studies

**Table 4:** comparison of WM specific and reasoning / contextually related dependent variables

| Author | Research design | Significance level of WM measure | Effect size of WM measure<br>0 =<small<br>1=small<br>2=medium<br>3=large |  | Most workplace relevant contextual outcome selected and reported here | Effect size of contextual measure<br>0 =<small<br>1=small<br>2=medium<br>3=large |  |
| --- | --- | --- | --- | --- | --- | --- | --- |
| Alloway & Warner, 2008 | Within contrasts repeated measures ANOVA | $F(1,18) = 6.08, p = .02$ | $d = 0.97$ | 3 | Reading and numerical test scores | Cannot calculate | |
| Ariës *et al.* 2014 study 1 | Within/Between ANOVA at each interval, group comparisons from final test presented here | $F(1,89) = 31.759, p = <.001$ | $r = .65$ | 3 | Reasoning abilities test scores (second interval) | $r = .13$ | 1 |
| Ariës *et al.*, 2014 | Within/between ANCOVA, | NS due to Bonferroni | *$d = 0.89^c$ | 3 | Reasoning abilities test | $r = .38$ | 2 |

<sup>c</sup> Where effect sizes are noted with \* they have been calculated by the author, not presented in the original paper

| study 2 | metacognitive training versus control at final test presented here | corrected $p$ value | | | scores within groups comparison | | |
| --- | --- | --- | --- | --- | --- | --- | --- |
| Chambers *et al.* 2008 | Within /between repeated measures ANOVA; Interval 2 control and intervention comparisons presented here | $F(1, 39) = 7.81, p=.01$ | $*d= 0.52$ | 2 | Mindfulness Awareness | $*d= 0.25$ | 1 |
| Craik *et al.*, 2007 | Between groups ANCOVA | $NS$ Means and SDs reported | $*d=0.1$ | 0 | Secondary Memory (Logical Stories) | $*d=0.66$ | 2 |
| Jha *et al.*, 2010 | Within groups (military trained) comparison paired samples $t$-test | Significant only with those reporting high practice, correlation | Cannot calculate for high practice groups only as M and | | Positive and Negative Affect respectively– NB only Intervention group Means | $*d=0.5$<br>$*d=0.5$ | 2<br>2 |
| | | between<br>practice<br>level and<br>WM<br>increase was<br>$r=.37$,<br>$p<.05$ | SDs<br>reported<br>for all<br>training<br>groups | | and SDs<br>provided | | |
| Miranda<br>*et al.*,<br>2011 | Between groups<br>ANCOVA<br>(baseline scores<br>as control<br>variable) post<br>intervention<br>scores<br>comparison<br>reported here | $F(1, 41) =$<br>5.558,<br>$p=.024$ | $\eta^2=.125$ | 2 | Attention<br>Vigilance<br>test | $\eta^2=.288$ | 3 |
| Moro *et al.*, 2012 | Within pre-post<br>(T1-T3) *t* -test<br>reported as<br>significant for<br>intervention<br>group A, effect<br>size calculated<br>from between<br>groups comparison at<br>T2 for<br>consistency,<br>where group B<br>act as a control<br>group | $t(14) = 2.48$,<br>$p = .027$ | *$d=0.8$ | 3 | Attention –<br>verbal span<br>test selected<br>as best work<br>related<br>measure,<br>again T2<br>between<br>groups comparison<br>selected | *$d=.84$ | 3 |
| Moro *et al.*, 2015 | Within pre-post<br>(T1-T2) *t*-test<br>reported as<br>significant for<br>intervention<br>group A, effect<br>size calculated<br>from between<br>groups<br>comparison at<br>T2 for<br>consistency,<br>where group B<br>act as a control<br>group | $t(14) = 2.3, p = .037$ | $*d = 1.28$ | 3 | Montreal<br>Overall<br>Cognitive<br>Assessment<br>was selected<br>as best work<br>related<br>measure,<br>again T2<br>between<br>groups<br>comparison<br>selected | $*d = 1.08$ | 3 |
| Zeidan, *et al.*, 2010 | Within/between<br>ANOVA,<br>session x group<br>reported here | $F(1, 47) = 1.26, p = .27$ | Cannot<br>calculate | | Fatigue | $*d = 0.7$ | 3 |
| Zylowska<br>*et al.*<br>2008 | Within groups<br>comparison only | $t(24) = 0.45$,<br>$p = .66$ | $*d = 0.1$ | 0 | ADHD<br>symptoms | $*d = 0.7$ | 2 |
| Average<br>effect<br>size |  |  |  | **2.1** |  |  | **2.7** |

Table 3 compares the WM studies, where the mean population ages ranged from 7.3 to 75 years; the statistical methods used were ANOVAs, ANCOVAs and *t*-tests. The studies used a variety of WM measures, all standardized and previously validated. Table 4 compares the WM effect sizes with more contextually based outcome effect sizes. These were determined as the most workplace relevant variable, all represented some form of higher order cognition (Hock, 2012) or emotional management and were consistent with the explicit training goals and context of the intervention, in some cases more so than WM. We are unable to calculate a reliable Hedges *g*, due to the small number of primary studies and inconsistent reporting of means and standard deviations. A crude but pragmatic effect size score was computed in order to compare effect sizes magnitude interpreted from the different measures (Cohen’s *d*, effect size *r* and partial eta squared) these are noted in Table 4 and an average computed for both the WM and contextual measure reported (Durlak, 2009).

Table 5 outlines the main elements of the comparison of the CIMO stages of analysis for the SE studies where the mean population ages ranged from 18 to 50 years; the statistical methods used were ANOVAs and *t*-tests, and two studies included non-parametric analyses, using Wilcoxon and Mann-Whitney tests. Many SE studies used a published General SE Scale (GSES, n-6), others used Teacher SE (n-2), specifically constructed scales (n-2), Academic SE (n-1) and Study Skills SE (n-1). One study did not specify the scales used (n-1, Tsai et al., 2011) but clarified that it was created with reference to the SCLT. One study included a GSES measure as well as a Job SE measure (McGonagle et al., 2014).

**Table 5.** . Data extracted from self-efficacy studies

RUNNING HEAD: REVIEW OF COACHING INTERVENTIONS FOR DYSLEXIA
| Author | Context |  |  | Interventions |  |  | Mechanisms | Outcomes |  |
| --- | --- | --- | --- | --- | --- | --- | --- | --- | --- |
| | Nationality, sample size, % female | Health / work / education / occupational group | Neurodeficit or identified WM deficit | Teaching / learning methods used | Time spent in intervention | Reliability of SE measure, $\alpha$ | Learning / psychological theories applied | Effect size of SE measure | |
| Bell, Raczynski and Horne (2010) | US, 50, NK | Working in education | NK | Group-based knowledge transfer and discussion | 7 sessions | .94 | SCLT - teacher efficacy | $d = 0.5148$ | Medium |
| Engin and Cam (2009) | Turkey, 22, 100% | Working in health | NK - but nursing up to 10% (Sander son-Mann & McCan dless,<br>2006) | Group-based knowledge transfer and discussion | 5 sessions | .81 | SCLT – SE, autonomy | $r = 0.88$ | Large |
| Franklin<br>and Doran<br>(2009) | Austra<br>lia,<br>52,<br>59% | Educati<br>on | NK | 2<br>worksh<br>ops<br>followe<br>d by 4<br>paired<br>peer<br>coachin<br>g<br>sessions | 9<br>hour<br>s | .86<br>Cited<br>in<br>Grant<br>&<br>Frankl<br>in<br>(2007<br>) | PAAL;<br>SCLT -<br>SE;<br>incremen<br>tal<br>implicit<br>person<br>theory | PA<br>AL<br>grou<br>p *d*<br>= -<br>1.21<br>;<br>Self-<br>reg<br>grou<br>p -<br>*d*=.1<br>.08 | Larg<br>e |
| McDowall<br>and<br>Butterwort<br>h (2014) | UK,<br>32,<br>75% | Educati<br>on<br>transitio<br>n to<br>work | NK | Group<br>coachin<br>g -<br>facilitat<br>ed<br>discussi<br>on | 1<br>sessi<br>on | .78 | Strengths<br>-based<br>coaching |  |  |
| McGonagl<br>e et al.<br>(2014) | USA,<br>59,<br>86% | Work | Chroni<br>c health<br>conditi | 1:1<br>phone<br>coachin | 6<br>sessi<br>ons | GSES<br>:.80-<br>.89 | SCLT –<br>SE, also<br>transacti | GSE<br>S: | Larg<br>e |
| | | | ons | g | of 1<br>hour | cited<br>in<br>Judge,<br>Erez,<br>Bono,<br>&<br>Thore<br>sen<br>(2003<br>)<br>Job<br>SE<br>.77-<br>.83<br>cited<br>in<br>Chen,<br>Godd<br>ard, &<br>Caspe<br>r,<br>(2004<br>) | onal and<br>conserva<br>tion of<br>resources<br>models<br>of stress | Parti<br>al<br>$\eta^2$<br>.18;<br>Job<br>SE:<br>Parti<br>al<br>$\eta^2$<br>.09 | Med<br>ium |
| McDowall | UK, | Work | NK | 1:1 | 1 | 0.83 | Apprecia | $\eta^2 =$ | Larg |
| , Freemann and Marshall (2014) | 54, 65% |  |  | coaching, two intervention conditions | session |  | active enquiry; feedback intervention theory | .24 | e |
| Reed, Kennett, Lewis and Lund-Lucas (2011) | Canada, 41, NK | Education | 20% | Seminar group with mixed info transfer and group discussion | NK - > 6 | 89<br>ited in Kenne tt & Reed (2009 ) | Learned resourcefulness | $d = .81$ | Large |
| Reif, de Vries, Petermann and Gorres (2013) | Germany, 234, 80% | Health | n/k | Seminar group with mixed info transfer and group discussion on | 8 x 90 mins | .76-.9 | 'Psycho-education' | $\eta^2 = .10$ | Medium |
| Stensrud, Gulbrandsen, Mjaaland, Skretting and Finset (2014) | Norway, 21, 29% | GPs at work | n/k | Role-play and debrief | 5 x 4 hours | 0.94 | SE | n/a | N/A |
| Style and Boniwell (2010) | UK, 93, NK | Experimental | n/k | Group discussion 1/3; peer coaching 1/3; self-reflection 1/3 | 6 sessions | .76-.9 | Positive psychology | $d = .55$ | Medium |
| Tsai et al. (2011) | Taiwan, 395, 98% | Working in health | NK - but nursing up to 10% (Sander son-Mann | Group training | 1.5 hours | 0.94 | Not stated at all | $r = .26$ | Small |
|  |  |  | &<br>McCan<br>dless,<br>2006) |  |  |  |  |  |  |
| Tschannen<br>-Moran<br>and<br>McMaster<br>(2009) | US,<br>93,<br>NK | Workin<br>g in<br>educati<br>on | n/k | Small<br>group<br>coachin<br>g; 1:1<br>coachin<br>g;<br>observa<br>tional<br>'live'<br>coachin<br>g | 5.75<br>hour<br>s | 0.9 | SCLT | $r =$<br>.24 | Smal<br>l |
| Watt,<br>Murphy,<br>Pascoe,<br>Scanlon,<br>and Gan<br>(2011) | Austra<br>lian,<br>118,<br>89% | Studyin<br>g<br>nursing | NK -<br>but<br>nursing<br>up to<br>10%<br>(Sander<br>son-Mann<br>&<br>McCan dless,<br>2006) | “Struct<br>ured<br>learning<br>progra<br>m” | 3<br>days | 0.69 | Weak<br>but<br>SCLT<br>related | $d =$<br>.87 | Larg<br>e |
| Zwerver,<br>Schellart,<br>Anema<br>and van<br>der Beek<br>(2013) | Holla<br>nd,<br>40,<br>50% | Workin<br>g as<br>doctors | n/k | Info<br>transfer,<br>role<br>play,<br>feedbac<br>k | n/k | .75-<br>.86 | Theory<br>of<br>planned<br>behavior,<br>attitude,<br>social<br>norms<br>and SE<br>model | $r =$<br>.33 | Med<br>ium |

## Synthesis of Findings

Whilst we outline the study findings for WM and SE separately, as the nature of these studies differed in terms of theoretical framing as well as the methods used, we had a common analytic strategy focusing on a direct comparison between ‘successful’ (i.e., an observed effect in the expected direction) and ‘unsuccessful’ interventions to isolate effective treatment mechanisms across studies. This approach led us to note the methodological shortcomings and lack of detail in the primary studies, which we note in each synthesis as appropriate, followed by an overall discussion in the final section of the paper.

## WM Synthesis

### Results for WM improvement

The effect sizes of the successful studies ranged from medium to large, with only one statistically significant study (Chambers et al. 2008) falling short of medium effect size correlation (*r* = .25). When mean data were presented, studies indicated an improvement of between a half and a whole standard deviation (Ariës et al., 2014; Chambers et al., 2008; Jha et al., 2010; Miranda, Presentacion, Siegenthaler & Jara, 2013). Moro et al. (2012, 2015) reported less than half a standard deviation improvement, but their sample experienced age-related cognitive decline and therefore have additional barriers when compared with dyslexia or working adults. Four studies reported non-significant results and the two effect sizes that were calculable for these were below the small range. The effect sizes for successful studies were similar to meta-analytic moderate aggregate effect sizes achieved for standard verbal WM through adaptive computerized WM training (*g* = .31: Melby-Lervåg & Hulme, 2012; *g* = .36, Weicker & Thöne-otto, 2015). We cannot discount that a sampling bias is present. We do, however, propose that the WM intervention effect is comparable for coaching interventions and computerized training, which of interest considering the differentiation in learner experience. Table 4 denotes a crudely computed average of medium effect size for the WM measure (2.1) indicating the bottom of the medium range, these calculations lack sophistication such as sample size weighting, yet the studies all reported relatively small sample sizes. We position the effect size interpretation as a trend indicating an avenue for further investigation, noteworthy mainly in its convergence with computerized WM training research but not reliable due to the limited extraction in this thesis.

### Intervention analysis

The interventions were typically a series of sessions that were conducted by either a professional in education/health or a trained meditation facilitator, although some studies additionally utilized peer learning. The number of sessions varied, with as many as 65 sessions delivered over one year (Alloway & Warner, 2008) yet as few as four sessions for two of the studies (Craik et al., 2007; Zeidan et al., 2010) who both notably reported no impact of their intervention on WM.

Comparing the quality of the studies was the most challenging, as the descriptions of the interventions varied considerably; we noted a tendency for researchers to very briefly describe the content of the intervention, leaving little opportunity for replication or analysis of the delivery. The theoretical components of the extracted studies were devoted to the WM-dependent aspects of learning, as opposed to the learning processes used during the intervention. Our initial review revealed a consistent picture in the successful studies: the activities described in the methods could be interpreted to represent the four critical elements of SCLT: (1) verbal persuasion (an introductory knowledge transfer in most cases); (2) role-modeling (either current/past case study discussion or active role play); (3) vicarious learning (group discussion) and (4) mastery (the opportunity to practice or rehearse in context, with recognition of success). In studies by Aries (2015, study 2, metacognition group only) Craik et al. (2007) and Zeidan et al. (2010), a lack of fidelity to SCLT resulted from an insufficient amount of time available to develop mastery and/or engage socially and these studies showed no improvement; however, Zylowska et al. (2008) provided sufficient time and discussion yet no improvement was noted in their study either. In analyzing this fourth study to identify the missing factor, we noted that WM was not a stated outcome of the intervention in Zylowska et al. (2008), and that specific practice of metacognition or stress management were not mandated. To summarize, we noted in addition to the general relevance of SE for dyslexic learners and employees presented in the introduction and expert panel contribution, interventions that supported the development of memory specific SE (through SCLT compliant activities) resulted in the improvement of WM, our finding has some support in the clinical literature (Valentijn et al., 2006). Our observed successful intervention protocol are in contrast to interventions involving the passive engagement of recipients in lessons involving knowledge transfer of memory strategies only (didactic teaching) or practice divorced from context (WM training games).

Further exploration of the how participants became engaged during the studies revealed a second key concept: ‘Metacognition’, or the development of self-awareness in thinking (Flavell, 1979), which was mentioned explicitly in four studies (Ariës et al., 2014 study 1 & 2; Moro et al., 2012, 2015) and by association with similar terms in two others (‘attentional control’, Chambers et al., 2008; ‘cognitive control’, Jha et al., 2010). Becoming aware of and deliberately manipulating thoughts to improve memory has some support in the literature on dyslexia (Gerber, 2012; Leather et al., 2011); the ‘Gerber-Leather model’ (de Beer et al., 2014) proposes the self-regulation of memory (WM) and positive reframing of the individual’s personal dyslexic experience as mediators of an improved sense of control, thereby influencing success in the workplace. Additional support is found in the clinical dementia research, in which ‘meta-memory’ (the ability to consciously be aware of and control mental memory tasks, such as visualizing a shopping list; Dixon & Hultsch, 1984) is improved through developing mental strategies and focusing on SE (Dunlosky, Bailey, & Hertzog, 2011; Jopp & Hertzog, 2007; Van der Elst, Van Boxtel, Van Breukelen, & Jolles, 2008). The extracted studies support these theories.

Two studies developed metacognition through mindfulness and meditation protocols, (Chambers et al., 2008; Jha et al., 2010) demonstrating an increase in WM concurrent with a decrease in negative emotions (negative affect and stress, respectively). Since *increases* in anxiety and stress are known to reduce WM capacity (Johnson, 2015; Otto et al., 2016), our synthesis indicates an argument for *reductions* of stress as a moderating variable; this again is further supported by dementia research (Kaszniak, 2011) and connects with our target population, since research indicates stress management as problematic for dyslexic employees (Doyle & McDowall, 2015). General mindfulness, unlike the targeted development of meta-memory, might not directly lead to improved WM but potentially acts via the mediator of reduced stress.

### Contextually-based outcome measures

The extracted context-based outcomes (Table 4) were heterogenous in nature but consistent in reporting improvements with medium effect sizes on average and appropriately significant *p* values for all studies, unlike the WM scores which included four unsuccessful interventions The crude computation aggregate effect size was 2.7, towards the top of the medium range and representative of a stronger effect than the WM measure. These results are divergent from results for contextual outcomes resulting from adaptive computerized WM training where small effect sizes are noted for far transfer measures such as verbal abilities or everyday life measures (respectively: *g* = .24: Melby-Lervåg & Hulme, 2012; *g* = .29, Weicker & Thöne-otto, 2015). In general published systematic reviews of computerized training (Weicker & Thöne-otto 2015; Melby-Lervåg & Hulme 2012; Dunning et al. 2013) report consistently stronger WM scores compared to weaker and in some cases unsuccessful effects for contextual measures, termed ‘far transfer’, since in the computerized paradigm they are further from the trained tasks, whereas in contextual, discursive coaching the WM measure is the far transfer. We therefore observe that the two divergent learning environments produce an opposite effect: where WM tasks are trained explicitly (as in computerized training) there is a medium effect size for highly related WM outcomes and consistently smaller effect for more contextually-based outcomes; where training is contextual (as presented here) there are more compelling effect sizes for the contextual outcome and marginally smaller, inconsistent yet also on average medium effect sizes for the WM outcome. Some contextual training appears to improve core WM without direct practice of WM tasks, as observed in Chambers (2008); Moro et al, (2012; More et al. 2015; Alloway & Warner (2008) and Jha etal. (2010); equally some WM computerized training improves contextual measures (Holmes et al. 2009). The outcomes appear to overlap, but the causal direction and transfer pathways between WM and contextual performance are neither clear nor reliable in either body of research.

## SE Synthesis

### Results for SE improvment

Three studies reported small effect sizes, and the remainder (eleven studies) reported medium to large effects. These effect sizes compare favorably to those measuring the ability of online training to improve SE, for example, which has generally shown smaller effects (see systematic review: Mccutcheon, Lohan, Traynor, & Martin, 2015). The studies tended to include other measures of mood and affect, and eight studies included measures of academic or work-related performance, mainly adherence to a new tool, skill or process, all of which were significantly improved as a result of the intervention. We observe that coaching is a reliable intervention for both improving SE and transferring learning to workplace abilities.

### Intervention analysis

The participants in the SE studies were typically adults who were engaged in learning related to their studies or work. The interventions tended to be delivered by a mix of professional educators and facilitators trained in a specific process (e.g., Franklin & Doran, 2009; McDowall et al., 2014), although some studies additionally utilized peer-to-peer coaching without providing adequate descriptions of the training provided to peer coaches (Franklin & Doran, 2009; McDowall & Butterworth, 2014; Style & Boniwell, 2010). As with the WM studies, and as expected for developing SE, the successful interventions involved all four elements of SCLT either overtly within the intervention structure or by allowing time for development of mastery before reassessment. In most cases, the participants’ SE was developed in relation to a clear and measurable learning outcome or goal related to their work or life, rather than directly targeting SE; this approach is congruent with Bandura’s (1986) original proposition. We thus observed a common trend to deliver interventions consistently with Goal Setting Theory (GST; Locke & Latham, 2002) in addition to SCLT. GST predicts that ‘goal clarity’ (GS1) focuses attention and inspires effort and persistence to achieve while creating the conditions for metacognition around the target behavior; this element was clearly adhered to in the extracted studies through the verbal persuasion element. However GST further proposes two other moderators for improvements in work performance: (GS2) SE for achieving the goals and (GS3) the commitments made to others in relation to the goals. The extracted studies’ methodologies reported sufficient attention to GS2 through mastery and rehearsal, but the results varied considerably regarding GS3. It was possible to infer fidelity to GS3 through the common occupational or educational contexts of each sample, with the exception of the study by Style and Boniwell (2010), yet this connection is tenuous. We propose that SE is not an isolated psychological construct and develops with positive, well contextualized and internalized goals.

In contrast to the results of the WM studies, where practise time was a key determinant of success, the time spent in interventions did not affect the level of significance or effect size of the SE studies; even those with a single intervention session reported a significant impact on SE (McDowall & Butterworth, 2014; McDowall et al., 2014). The interventions were typically shorter than those in the WM papers, with the longest being five sessions of four hours; however, poor descriptions were common again, one paper not reporting the intervention length at all (Zwerver et al., 2013). Neither time nor the use of trained vs. peer coaches could be identified as a determinant of effect size range. The use of general vs. context-specific scales did not result in any significant patterns.

The single extracted study in which no improvement was found at all (Stensrud, 2014) reassessed SE at the immediate end of the program, before the participants could practice skills in their own setting (i.e., to develop mastery), and thus the results may have reflected a methodological artifact caused by testing before mastery had been attained. Indeed, Tsai et al. (2011) observed that SE decreased in the period immediately after the intervention before recovering to an increase from baseline after three months, indicating again that time to practice was needed to obtain mastery. In the single session interventions, mastery was incorporated by asking the participants to recall and explore incidences of previous mastery (McDowall & Butterworth, 2014; McDowall et al., 2014). Therefore, as with the WM studies, the practice/rehearsal/mastery element is highlighted as a key variable in successful interventions. The cohesion between learning goals and socially interactive development of SE facilitated consistently successful outcomes.

## Discussion

The results of our search demonstrated that an insufficient number of studies have evaluated learning or coaching activities for dyslexia, or other relevant context with robust and appropriate study designs, as published research in general provides. Given the relatively small number of studies reviewed overall, the implications for research and practice outlined below are limited and should be interpreted with caution and our propositions viewed as tentative hypotheses for further testing in studies with an appropriately sampled, longitudinal design.

Regarding our primary research question,: ‘to what extent, and under what conditions, can face-to-face (C) learning interventions (I) improve WM (MO_1_) and SE (MO_2_)’, fidelity to SCLT and GST protocols provided consisted evidence of the effectiveness of coaching to improve SE, as is practiced in UK disability accommodation coaching programs and thereby supporting practice for this variable. Evidence that coaching improves WM was less consistent and also apparently contingent on fidelity to SCLT (in particular mastery experiences) and development of self-awareness / stress management through metacognitive practice. The salient variables are described in Fig. 2, which depicts proposed intervention protocol that can be applied to facilitate disability accommodation through coaching dyslexic adults in the workplace. We should note with caution that an active goal setting component was inferred through the intervention description rather than through explicit statements in the studies with the exception of McDowall et al. (2014). The lack of detail in intervention protocols and reliance on our interpretation of key phrases such as ‘discussion’ and ‘reflection’ to infer dialetic principles was a limitation of our synthesis. We note that grey literature was not included in this extraction and, though this was a deliberate attempt to isolate research based evidence regarding psychological variables to refute or vindicate contemporary practitioner guidance (McLoughlin & Leather, 2013; Moody, 2010) systematic review of practitioner evidence may yet strengthen our findings.

**Figure 2.**
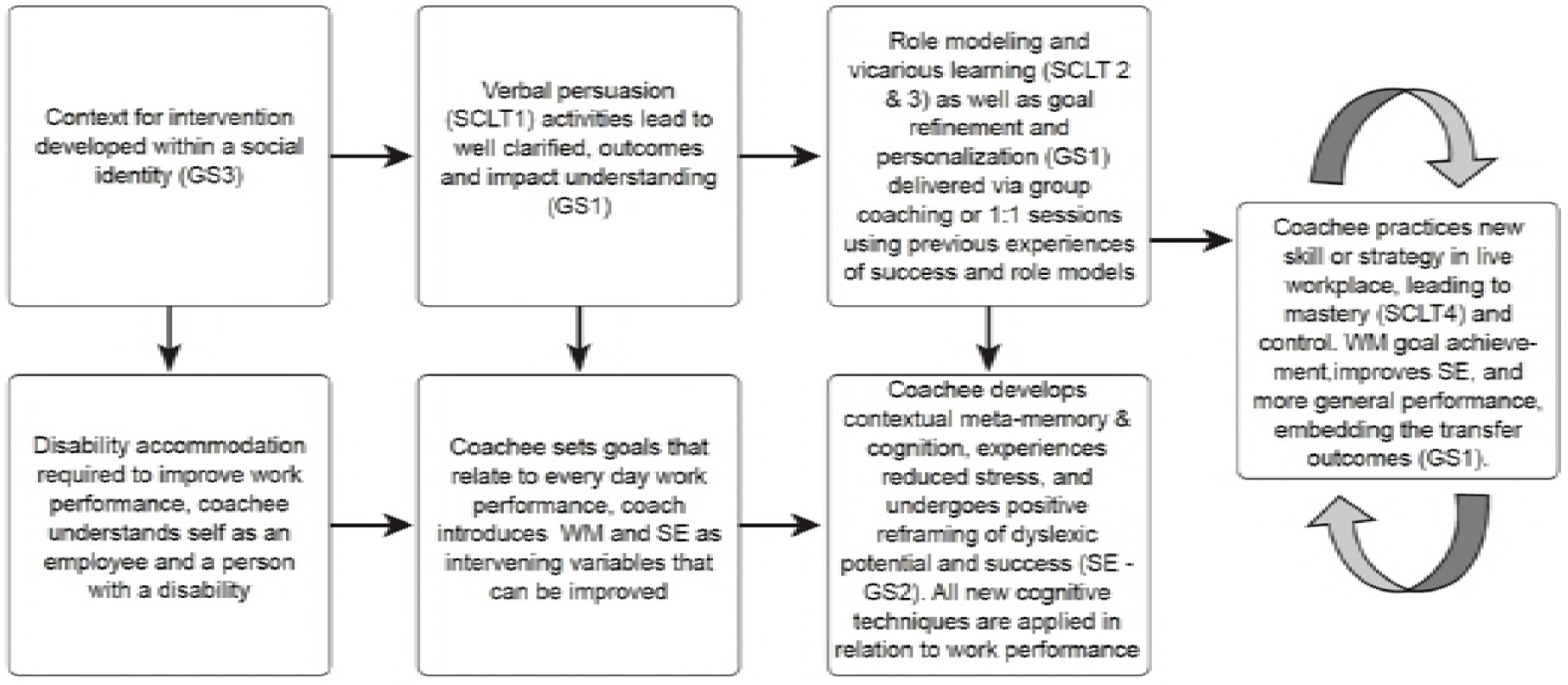
Hypothetical use of SCLT, metacognitive development and GST to improve SE & WM through face-to-face learning interventions.

The consistency between the WM and SE studies regarding the application of the SCLT was surprising and clearly showed that any new skill, including development and management of cognitive function such as WM, needs to be facilitated while considering the individual learner’s social and metacognitive experience; additionally, particular attention should be devoted to practice opportunities when developing a new skill. WM skills, which are reported to be of highest concern to our target population, cannot simply be ‘taught’; learners must develop them for themselves, with support and reflection on mastery opportunities (Valentijn et al., 2006).

## Relevance to Target Population

It was a significant limitation of our narrative review that there is no direct evidence on workplace disability accommodations through coaching based interventions to adult dyslexics, hence we broadened our inclusion criteria to include studies undertaken with other samples including ADHD, clinical and healthy, students and non-dyslexic adults. We now address this feature of the extraction by directly analyzing any differences in population or intervention protocols that would inhibit transfer to dyslexia accommodation coaching, in order to assess whether, in principle, the findings of our synthesis could be applied to our specific context.

### Population

The search protocol included a broad age range (from 7 to 75 years); however, our target population was working adults. Successful studies included all age ranges, and combined, the results demonstrated that in principle, improvement in WM was possible regardless of age. All SE studies were based on working age adults and were therefore directly comparable, but in particular, we note McGonagle et al. (2014), who provided coaching as a disability accommodation to a sample of working adults. The samples of individuals with dyslexia were not extracted directly, but when dyslexia or other developmental learning difficulties were included in the samples, improvements did occur (Alloway & Warn, 2008; Ariës et al., 2014; Miranda et al., 2013). Gender was well balanced between the studies and did not affect the results. The population comparison did not clearly support applicability to adults with dyslexia, but the authors conclude that there is potential for generalizability, thereby providing a premise for further direct evaluation research.

### Intervention protocol

The current practice in dyslexia coaching is to provide an average of 4-5 sessions delivered on a one-to-one basis (Doyle & McDowall, 2015). The extracted studies that matched this structure were broadly successful, indicating that typical dyslexia coaching protocols may potentially meet the requirements of facilitating WM and SE improvement; however, the quality of coaching and adherence to SCLT and GST may be as important as sufficient intervention time.

One major difference between the extracted studies and the current models of dyslexia coaching is the use of group versus one-to-one training. Some studies included one-to-one elements (McDowall & Butterworth, 2014; McDowall et al., 2014; McGonagle et al., 2014), yet all intervention protocols except one (McGonagle et al., 2014) reported some element of group discussion, peer coaching or coaching triads. The one-to-one intervention achieved a positive result; however, a greater understanding of the group dynamics in WM and SE outcomes is needed. It is possible that a dyslexic population might benefit further from group coaching, but no studies have compared the impact of group coaching with one-on-one protocols.

In summary, though our extracted studies did not match our target population, we have addressed the fundamental challenge of reviewing diverse research with potential relevance, to provide a narrative, conceptual framework justifying further, primary studies. In doing so, we avoid ‘reinventing the wheel’ and reduce the need to conduct primary, exploratory studies on the viability of coaching as a mechanism to effect WM and SE for dyslexia, (Briner & Rousseau, 2011) rather we can build on existing proof of principle in this research and progress the hypothetical pathway for change straight to intervention evaluation for dyslexic adults.

## Implications for WM research

Whilst initially not a core focus for our investigation, and limited by the small number of heterogenous samples, the findings from this systematic review provide insight on WM research in general, when compared with systematic reviews of WM computerized training (Melby-Lervåg et al., 2016; Weicker & Thöne-otto, 2015). We suggest that more work is required to understand the dynamic relationship between WM and contextual outcomes. While previous WM training research has been honing the computerized training environment, the effect on contextually relevant ‘far transfer’ measures has not been forthcoming, which questions the role of WM as foundation for developing complex reasoning skills, contrary to previous conceptual work (Bailey, 2007; Mantyla & Carelli, 2006; Thorell et al., 2009; Varvara, Varuzza, Sorrentino, Vicari, & Menghini, 2014) and concurrent with previous critiques that WM, or at least the measurement thereof, lacks ecological validity (Chaytor, Schmitter-Edgecombe, & Burr, 2006; Van der Elst et al., 2008).

The studies presented in this paper suggest that instead, directly targeting complex reasoning skills such as self-regulated attention, metacognition, and behavioral self-awareness can in fact improve WM, achieving moderate success in both outcome measures, even when WM itself was not a directly expressed goal of the intervention (Chambers et al., 2008; Jha et al., 2010). We note the limitation of the primary papers to provide sufficient data for refined effect size aggregation, yet the trend warrants more research at least. It is not clear how the improved complex reasoning tasks lead to improved WM, nor whether direct targeting of WM will ever lead to consistent far transfer effects; we need to develop our understanding of the other ‘active ingredients’ in learning, comprehension and applied cognitive skills. Potential pathways for future research include (1) the reduction of stress, (2) the role of developing SE in targeted tasks and (3) the ability of metacognitive awareness to be applied to multiple areas of cognition once learned in one area. We propose that the incorporation of metacognition and SCLT in computerized WM training research may add depth to further our understanding of effective memory processes and strategies as well as outcomes. Additionally, GST incorporated in to the verbal persuasion element of SCLT, inclusion of peer discussion or verbal debriefing, contextualized mastery opportunities with dialectical reflection might provide rich areas for experimental investigation.

## Implications for practice

Our initial scoping process identified that SE and WM improvements are key ‘active ingredients’ for dyslexic people at work, our synthesis indicates that coaching is a potential vehicle for facilitating improvements, as is common in practice for dyslexia (and indeed other ‘hidden’ disabilities) in workplace disability accommodation. Improvements were facilitated by a SCLT compliant protocol, where time spent in intervention was less important than the quality of the intervention and the opportunities for the development of mastery. Contextualised, work-related outcomes were positively impacted by the coaching interventions, at times irrespective of WM improvement, but not irrespective of SE improvement for which clear evidence in support of coaching interventions was found. In order to apply these findings to practice, psychologists should ensure the inclusion of Goal Setting Theory and SCLT based interventions when making recommendations for disability accommodation in workplaces, bearing in mind the potentially flawed generalizability from group to 1:1 contexts and the need for process evaluation in context.

## Conclusion

This review demonstrates an urgent need for future robust field research to investigate the effectiveness of coaching and learning interventions delivered as disability accommodation in occupational contexts with distinct focus on transfer of learning into workplace performance. Whilst we note the small number and variable quality of primary studies, the results for WM question appear to question the effectiveness of current research interventions, insofar as the as evidence for real world outcomes appears limited (Chaytor et al., 2006; Van der Elst et al., 2008). In the context of dyslexia, which was the impetus for the current paper, we argue for greater contextualization of any interventions, which should pragmatically combine neuropsychological, behavioural and social variables into a biopsychosocial model of dyslexia to build on the existing, comparatively robust, evidence regarding literacy support and the development of SE. The next step is to translate these findings into controlled, longitudinal, intervention evaluations to further elucidate the role of SCLT, GST, metacognition and stress management in disability accommodation for dyslexia.

Those emboldened were included in the synthesis.

Contributors
The first author conducted the expert consultation, searches, extractions and quality check and synthesis. The second author second checked all searches, extractions, quality check and synthesis. The first author wrote the initial draft, the second contributed significant edits and re-drafted large portions of the paper. Both authors edited the document until complete.

